# Towards interpretable molecular and spatial analysis of the tumor microenvironment from digital histopathology images with HistoTME-v2

**DOI:** 10.1101/2025.06.11.658673

**Authors:** Sushant Patkar, Timothy R. Rosean, Palak Patel, Stephanie Harmon, Peter Choyke, Tamara Jamaspishvili, Baris Turkbey

**Affiliations:** Artificial Intelligence Resource (AIR), National Cancer Institute, National Institutes of Health, Bethesda, MD, USA; Department of Pathology and Laboratory of Medicine, SUNY Upstate Medical University, Syracuse, NY, USA

## Abstract

The tumor microenvironment (TME) is a critical focus for biomarker discovery and therapeutic targeting in cancer. However, widespread clinical adoption of TME profiling is hindered by the high cost and technical complexity of current platforms such as spatial transcriptomics and proteomics. Artificial Intelligence (AI)-based analysis of the TME from routine Hematoxylin & Eosin (H&E)-stained pathology slides presents a promising alternative. Yet, most existing deep learning approaches depend on extensive high-quality single-cell or patch-level annotations, which are labor-intensive and costly to generate. To address these limitations, we previously introduced HistoTME, a weakly supervised deep learning framework that predicts the activity of cell type-specific transcriptomic signatures directly from whole slide H&E images of non-small cell lung cancer. This enables rapid, high throughput analysis of the TME composition from whole slide H&E images (WSI) without the need for segmenting and classifying individual cells. In this work, we present HistoTME-v2, a pan-cancer extension of HistoTME, applied across 25 solid tumor types, substantially broadening the scope of prior efforts. HistoTME-v2 demonstrates high accuracy for predicting cell type-specific transcriptomic signature activity from H&E images, achieving a median Pearson correlation of 0.61 with ground truth measurements in internal cross-validation on The Cancer Genome Atlas (TCGA), encompassing 7,586 WSIs, 6,901 patients, and 24 cancer types, and a median Pearson correlation of 0.53 on external validation datasets spanning 5,657 WSIs, 1,775 patients and 9 cancer types. Furthermore, HistoTME- v2 resolves the spatial distribution of key immune and stromal cell types, exhibiting strong spatial concordance with single-cell measurements derived from multiplex imaging (CODEX, IHC) as well as Visium spatial transcriptomics, spanning 259 WSI, 154 patients, and 7 cancer types. Overall, across both bulk and spatial settings, HistoTME-v2 significantly outperforms baselines, positioning it as a robust, interpretable and cost-efficient tool for TME profiling and advancing the integration of spatial biology into routine pathology workflows.

## Introduction

The tumor microenvironment (TME) is a complex ecosystem of interacting cell types that collectively influence tumor progression, invasiveness, and therapeutic response^1,2^. Recent advances in in situ molecular profiling—such as spatial transcriptomics and proteomics— have enabled high-dimensional, spatially resolved analysis of the TME, revealing novel disease mechanisms and therapeutic targets^3–9^. However, translating these rich biological insights into clinically actionable biomarkers remains challenging due to high instrumentation costs, technical hurdles and the need for specialized expertise in multimodal data integration^10–13^.

In contrast, hematoxylin and eosin (H&E)-stained pathology slides are widely accessible in clinical settings. Artificial intelligence (AI)-driven systems, particularly deep learning models such as HoverNet^14^, CellVIT^15^ and others^16^ have shown promise in automated segmentation and classification of key cell types and tissue structures from digitized H&E images, offering a cost-efficient alternative to advanced molecular profiling. However, their performance depends heavily on access to large, high-quality, single-cell annotated datasets across diverse tissue types—resources that are labor-intensive to generate and subject to inter- observer variability^17–19^.

As an alternative, several studies have explored generative AI or contrastive learning-based approaches, which are designed to impute the spatial expression of various cell type markers directly from H&E images, leveraging paired H&E, multiplex imaging or spatial transcriptomics datasets for multimodal training^20–25^. While these methods show promise, they are often sensitive to variability in staining protocols across different centers^24,26^. Furthermore, acquiring large high-quality, spatially registered datasets remains a major challenge due to sectioning-induced tissue heterogeneity, deformation artifacts and the high data generation and storage costs^12,27^. Spatial transcriptomics datasets additionally face challenges of frequent gene dropouts and site-specific batch effects^28,29^. Collectively, these issues significantly constrain the quality and diversity of multimodal datasets available for training. As a result, most models are trained on a limited number of samples from a single cancer type, which restricts their generalizability across broader oncologic contexts^30,31^.

To overcome these hurdles, we recently introduced HistoTME^32^, a novel weakly supervised deep learning framework designed for rapid, high-throughput molecular profiling of the TME from routine H&E-stained slides. Conceptually, instead of segmenting and classifying individual nuclei or generating virtual stains, we reframe the problem as a weakly supervised regression task. Given a whole-slide H&E image as input, we predict enrichment scores for specific cell types or pathways within the whole tissue. This strategy allows HistoTME to learn subtle morphological patterns associated with the presence of specific cell types or activity of specific biological processes in an unbiased manner using widely available H&E images and paired bulk transcriptomics datasets for multimodal training. Moreover, by linking complex morphological features to biologically interpretable bulk transcriptomic signatures, our model has the potential to reveal novel imaging biomarkers that are often missed by trained pathologists and remain inaccessible without developing specialized molecular stains^30^.

Building on this foundation, we introduce *HistoTME-v2*, which scales HistoTME to 25 distinct cancer types. HistoTME-v2 has been trained in a pan-cancer fashion on 7,586 whole-slide H&E images paired with bulk transcriptomic profiles from The Cancer Genome Atlas (TCGA)^33^ and externally validated on 5,657 paired whole slide H&E images and transcriptomics samples from CPTAC^34^ and other institutional cohorts^35,36^. HistoTME-v2 accurately predicts activity of 29 cell type-specific and pathway-level gene expression signatures^37^ from H&E images, achieving on average 53% improvement in Pearson correlation over the baseline, SEQUOIA^38^, a recent state-of-the-art pan-cancer gene expression prediction method.

Importantly, HistoTME-v2 enables spatially resolved predictions of key immune and stromal cell types within the TME, demonstrating superior generalizability relative to baseline spatial transcriptomics prediction methods, when benchmarked against Visium, multiplex IHC and CODEX datasets. Collectively, these results establish HistoTME-v2 as a robust, interpretable, and cost-effective platform for high-dimensional spatial analysis of the TME, further advancing biomarker discovery efforts from standard histopathology images.

## Results

### Overview of HistoTME-v2

HistoTME-v2 is a pan-cancer extension of our recently developed HistoTME model, which predicts 29 cell type specific and pathway-level gene expression signatures^37^ directly from H&E-stained whole-slide images (**Figure 1A**)^32^. These signatures serve as biomarkers for well-known tumor infiltrating immune and stromal cell populations (e.g., T cells, B cells, Macrophages, Fibroblasts and Endothelial cells) as well as key immunological and oncogenic processes active within the TME (e.g., antigen presentation, expression of immune checkpoints, cytokine/chemokine signaling, epithelial to mesenchymal transition, cell proliferation and matrix remodeling). This allows us to comprehensively characterize the TME composition directly from H&E images without relying on computationally intensive single cell segmentation and classification or sophisticated staining protocols.

**Figure 1:**
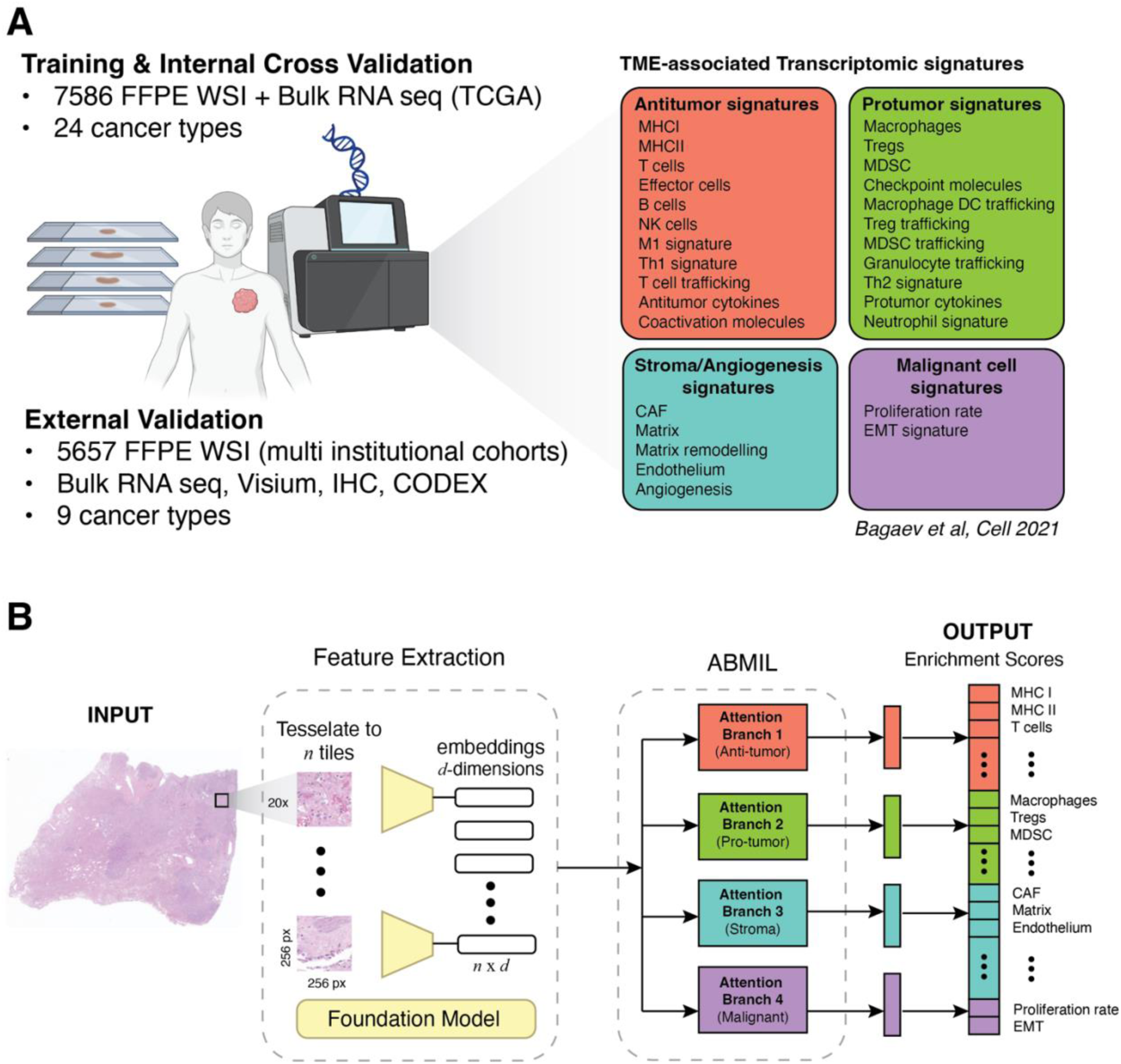
Overview of the HistoTME-v2 training pipeline and validation cohorts. **(A)** HistoTME-v2 is a pan- cancer extension of the HistoTME deep learning framework^32^, designed to predict cell type– and pathway- level gene set enrichment scores directly from whole slide H&E images (WSIs). The model was trained using 7,586 paired WSIs and bulk tumor transcriptomic profiles from The Cancer Genome Atlas (TCGA)^33^ across 24 solid tumor types, using 5-fold cross-validation with 20% of each fold held out for internal validation. External validation was performed on 5,657 FFPE WSIs spanning 9 cancer types from multi-institutional cohorts including CPTAC^34^, HEST1K^48^, NCI Prostate Cancer^36^, Fred Hutch High Grade Ovarian Cancer (HGSOC)^35^, SUNY Upstate Non-Small Cell Lung Cancer (NSCLC)^32^, and Colorectal Cancer (CRC)-FFPE-CODEX cohorts^45^. CPTAC, NCI, and Fred Hutch cohorts included matched bulk RNA-seq data, enabling evaluation of transcriptomic signature prediction performance at the bulk level. SUNY Upstate, CRC, and HEST1K cohorts provided spatially resolved data—including multiplex IHC, CODEX, and Visium spatial transcriptomics—to assess model generalizability at bulk, regional, and tile-level spatial resolutions. This figure panel was generated with the help of biorender (https://www.biorender.com/). **(B)** The bottom panel illustrates the attention-based multiple instance learning (ABMIL) pipeline of HistoTME-v2, which operates on tessellated 256×256 pixel tiles derived from WSIs at 20x magnification.

The HistoTME-v2 pipeline begins by tessellating one or more whole-slide H&E images (WSI) per patient into non-overlapping tiles of size 256x256 pixels at 20x magnification (∼112x112 microns). Each tile is then converted into semantically rich feature embeddings using digital pathology foundation models. In our training experiments for HistoTME-v2, we explored feature representations from six different state-of-the-art digital pathology foundation models (UNI^39^, UNI2^39^, Virchow^40^, Virchow2^41^, Gigapath^42^, and Hoptimus0^43^). After feature extraction, HistoTME-v2 utilizes an attention-based multiple instance learning (ABMIL) model^44^ to dynamically aggregate all tile-level feature embeddings into a single bag representation. This bag representation is fed to a simple multilayer perceptron (MLP) to predict normalized gene set enrichment scores corresponding to 29 distinct cell types and pathways. A key characteristic of our deep learning architecture is that it utilizes shared attention branches for predicting the activity of established functionally correlated gene sets defined previously as follows: anti-tumor, pro-tumor, stroma/angiogenesis and malignant cell signatures^37^. In our preliminary work with HistoTME, we showed that this modular design encourages feature sharing across closely related signatures, thereby enhancing model generalizability^32^ (**Figure 1B**). Furthermore, leveraging ABMIL provides added flexibility of analyzing tissue samples of varying sizes and scales ranging from small TMA cores or needle biopsies to large surgical tumor sections.

### HistoTME-v2 accurately predicts TME-associated transcriptomic signature activity across multiple cancer types

HistoTME-v2 was trained in a 5-fold cross-validation fashion utilizing 7,586 tumor samples from 6901 patients in The Cancer Genome Atlas (TCGA; n=24 cancer types), with paired formalin-fixed paraffin-embedded (FFPE) H&E slides and bulk RNA-sequencing data. In each fold, 80% of the pan-cancer patient cohort was utilized for training and 20% was held out for validation. To measure accuracy, we quantified the Pearson’s correlation between HistoTME-v2-predicted and ground truth enrichment scores in each cancer type. Our cross-validation experiments revealed that an ensemble of foundation models, in general, yielded more accurate predictions compared to any single foundation model, achieving, on average, a high median Pearson correlation of 0.61 across signatures (**Figure 2A**). The most accurate predictions were observed in Thyroid Carcinoma (TCGA-THCA) samples with a median Pearson correlation of 0.72 across signatures, whereas the least accurate predictions were observed in Uterine Carcinosarcoma samples (TCGA-UCS) with a median Pearson correlation of 0.43 across signatures. Interestingly, the accuracy in most cancer types was consistently high despite the skewed cancer type representation in the TCGA pan-cancer cohort, underscoring its ability to learn generalizable representations of TME biology across histologically diverse tissues. The ensemble model was chosen as the final model for all subsequent external validations.

**Figure 2:**
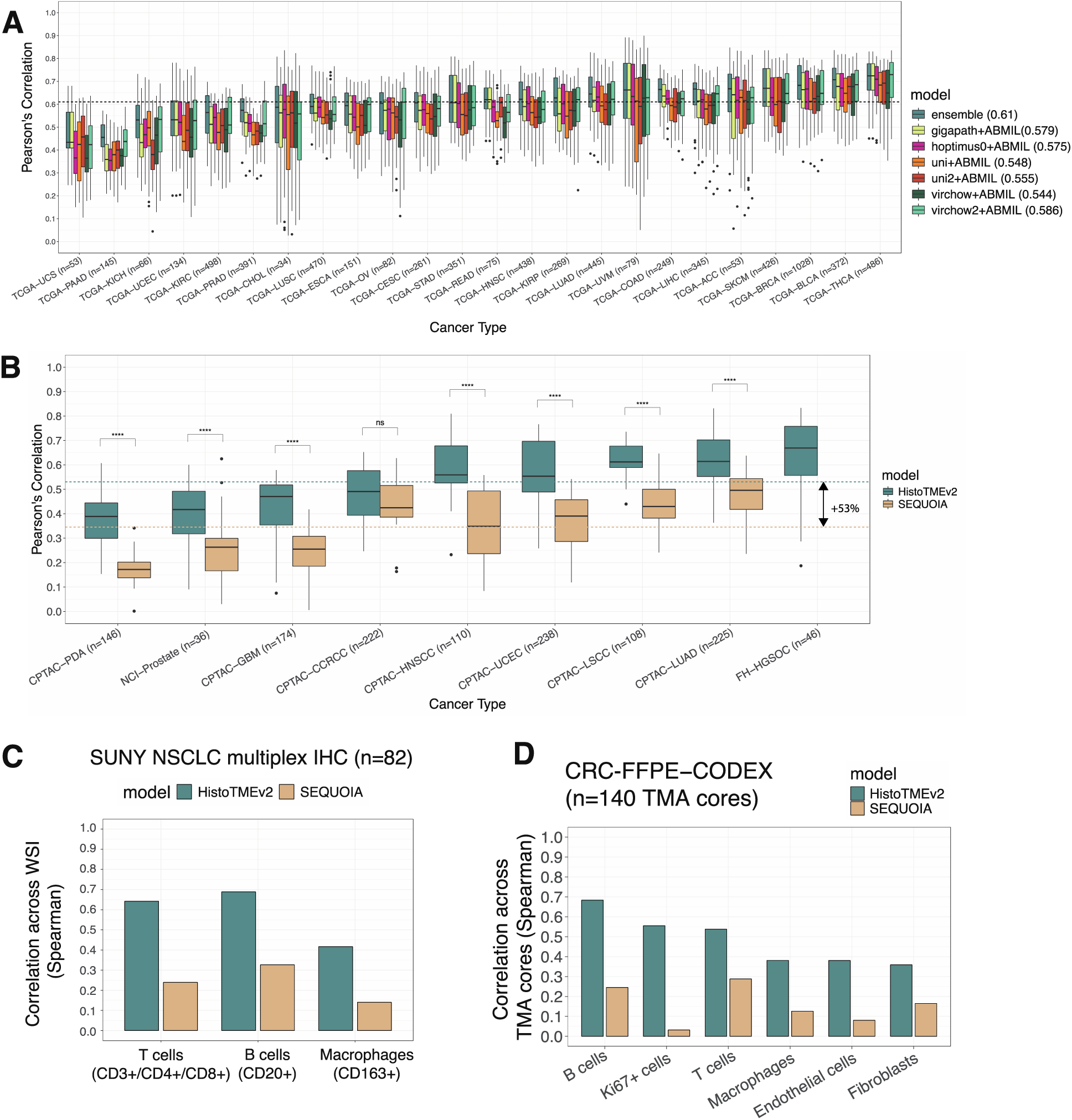
Pan-cancer internal and external validation of HistoTME-v2. **(A)** Boxplots showing the distribution of 5-fold cross-validation Pearson correlations for 29 predicted TME- related gene expression signatures across 24 cancer types in TCGA^33^, using matched FFPE H&E images and bulk RNA-seq data. The final model selected for external validation is an ensemble of these cross-validated models, achieving, on average, a median Pearson correlation of 0.61 across cancer types (shown as a black dashed line). The average median correlations for all other model variants are shown in the legend. Cohort number for each cancer type are reported in parentheses. Cancer types are sorted in increasing order of performance of the ensemble model from left to right. **(B)** External validation performance of HistoTME-v2 ensemble model on nine independent cohorts with matched FFPE H&E and bulk transcriptomic data, including CPTAC^34^, NCI Prostate (NCT02430480)^36^, and the Fred Hutch High-Grade Serous Ovarian Cancer (FH-HGSOC) cohort^35^. HistoTME-v2 is benchmarked against SEQUOIA^38^, that predicts whole-transcriptome expression from H&E images, from which signature scores were derived post hoc. SEQUOIA is trained separately on data for each cancer type, unlike HistoTME-v2. Predictions of SEQUOIA for the Fred Huch High Grade Serous Ovarian Cancer (HGSOC) cohort are missing due to lack of an ovarian cancer-specific pretrained SEQUOIA model. Cohort number for each cancer type are reported in parentheses. Green dashed line represents the average median Pearson correlation across signatures and cancer types for HistoTME-v2. The beige colored line represents the average of the median Pearson correlation across signatures and cancer types for SEQUOIA. **(C)** Comparison of HistoTME-v2 and SEQUOIA in predicting immune cell abundances in surgical NSCLC specimens from the SUNY Upstate NSCLC cohort^32^. Predicted enrichment scores for T cells, B cells, and macrophages were correlated with experimentally measured single-cell fractions (% marker positive cells) from serial multiplex IHC sections. **(D)** Region-level (∼960×720 µm) correlation of HistoTME-v2- and SEQUOIA-predicted enrichment scores with ground truth cell type fractions (T cells, B cells, macrophages, endothelial cells, fibroblasts, and Ki67+ cells) derived from CODEX imaging across 140 tissue microarray cores from the tumor invasive front of colorectal cancer patients (CRC-FFPE-CODEX cohort^45^).

Having trained HistoTME-v2 in a pan cancer fashion on TCGA, we next assessed its ability to generalize to independent multi-institutional tumor datasets (CPTAC, NCI-Prostate and Fred Huch HGSOC cohorts), spanning 5657 WSI, 1775 patients and 9 cancer types. For benchmarking, we compared HistoTME-v2 to a strong baseline: SEQUOIA^38^, a recently established state-of-the-art deep learning framework that predicts bulk gene expression from H&E slides of 16 tumor types. We computed gene set enrichment scores for each signature from SEQUOIA’s predicted transcriptomes, representing an *indirect* inference approach. In contrast, HistoTME-v2 learns to *directly* predict enrichment scores of curated gene signatures without constructing the entire transcriptome. HistoTME-v2 significantly outperformed SEQUOIA in predicting transcriptomic signature activity directly from H&E images, with an average improvement of 53% in Pearson correlations (**Figure 2B**). These results highlight the enhanced robustness of HistoTME-v2 in directly modeling transcriptomic signature activity over baseline H&E-based gene expression prediction methods.

### HistoTME-v2 captures immune and stromal cell composition at single-cell resolution

To further validate HistoTME-v2’s predictions at single-cell resolution, we assessed the concordance between its predicted cell type–specific enrichment scores and experimentally measured single-cell abundances derived from matched multiplex immunohistochemistry (IHC) and CODEX imaging datasets. These analyses focused on cell types for which both transcriptomic signature scores and corresponding protein markers were available, enabling direct one-to-one comparisons.

First, we applied HistoTME-v2 to an institutional cohort of 82 NSCLC surgical tumor specimens from SUNY Upstate University (SUNY Upstate Cohort^32^). Each tumor from this cohort had serially cut H&E and multiplex IHC sections available for T cells (CD4+ and CD8+), B cells (CD20+), and macrophages (CD163+). Single cell fractions of each cell type were quantified from corresponding IHC stained sections using the QuPath positive cell detection pipeline (See Methods). Overall, we observed strong positive correlations between HistoTME-v2-predicted enrichment scores for T cells, B cells and Macrophages and experimentally measured single cell fractions of the same cell types (**Figure 2C**, Spearman’s rho: 0.65 for T cells, 0.72 for B cells, 0.42 for Macrophages). In comparison, enrichment scores derived from SEQUOIA’s bulk gene expression predictions showed substantially weaker correlations (**Figure 2C**; Spearman’s rho: 0.24 for T cells, 0.38 for B cells, 0.14 for Macrophages).

Next, we assessed whether HistoTME-v2’s predicted enrichment scores align with single cell fractions derived from CODEX imaging. To do this, we utilized a publicly available dataset consisting of paired H&E and multiplex CODEX images from spatially distinct regions of the colorectal cancer TME (CRC-FFPE-CODEX cohort^45^; See Methods). In this dataset, we specifically focus on analysis of T cells, B cells, macrophages, fibroblasts, endothelial cells, and proliferating (Ki67+) cells within each tissue microarray (TMA) core, as both HistoTME- v2-predicted signature scores as well as ground truth single cell abundances derived from CODEX imaging markers were available for head-to-head comparison. We benchmarked HistoTME-v2-derived enrichment scores against SEQUOIA-derived enrichment scores, using the Spearman correlation metric. HistoTME-v2 outperformed SEQUOIA in capturing regional cell type-specific abundances across TMA cores. Specifically, HistoTME-v2’s enrichment scores showed stronger positive correlations with single cell abundances of T cells (ρ = 0.54 vs. 0.29), B cells (ρ = 0.68 vs. 0.25), Ki67⁺ cells (ρ = 0.56 vs. 0.12), macrophages (ρ = 0.38 vs. 0.13), fibroblasts (ρ = 0.36 vs. 0.17), and endothelial cells (ρ = 0.38 vs. 0.08), when compared to SEQUOIA-derived scores for the same cell types (**Figure 2D**).

Collectively, these findings demonstrate HistoTME-v2’s enhanced ability to estimate the TME composition from routine H&E images compared to existing state of the art bulk gene expression prediction methods.

### Benchmarking Spatial Prediction Performance of HistoTME-v2

Although HistoTME-v2 was designed to predict bulk TME composition, we next evaluated its capacity to resolve spatial heterogeneity within the TME. To this end, we conducted the following benchmarking experiments.

First, we investigated HistoTME-v2’s ability to map the spatial distribution of T cells, B cells, and macrophages at the individual tile-level (∼112 microns) in the SUNY Upstate NSCLC Cohort due to availability of corresponding spatially registered ground truth multiplex IHC stains. To generate spatially resolved predictions for each cell type, we applied the pretrained HistoTME-v2 model in a sliding window fashion on each WSI using a 3×3 tile window size (see **Figure 3A**). This design choice enables fine-grained spatial inference without introducing any modifications to the existing model architecture (See **Figure 3B**). For benchmarking, we compared HistoTME-v2 against two established spatial prediction methods: SEQUOIA, which infers spatial gene expression from H&E images using a similar sliding window approach, and DeepSpot ^46^, a deep learning model trained to predict spatial transcriptomes from H&E images using paired FFPE-H&E and Visium spatial transcriptomic datasets from HEST-1K and the University Hospital of Zurich. Since both SEQUOIA and DeepSpot produce spatial transcriptomic predictions, cell type enrichment scores were derived post hoc from these outputs for head-to-head comparison with HistoTME-v2 (see Methods).

**Figure 3:**
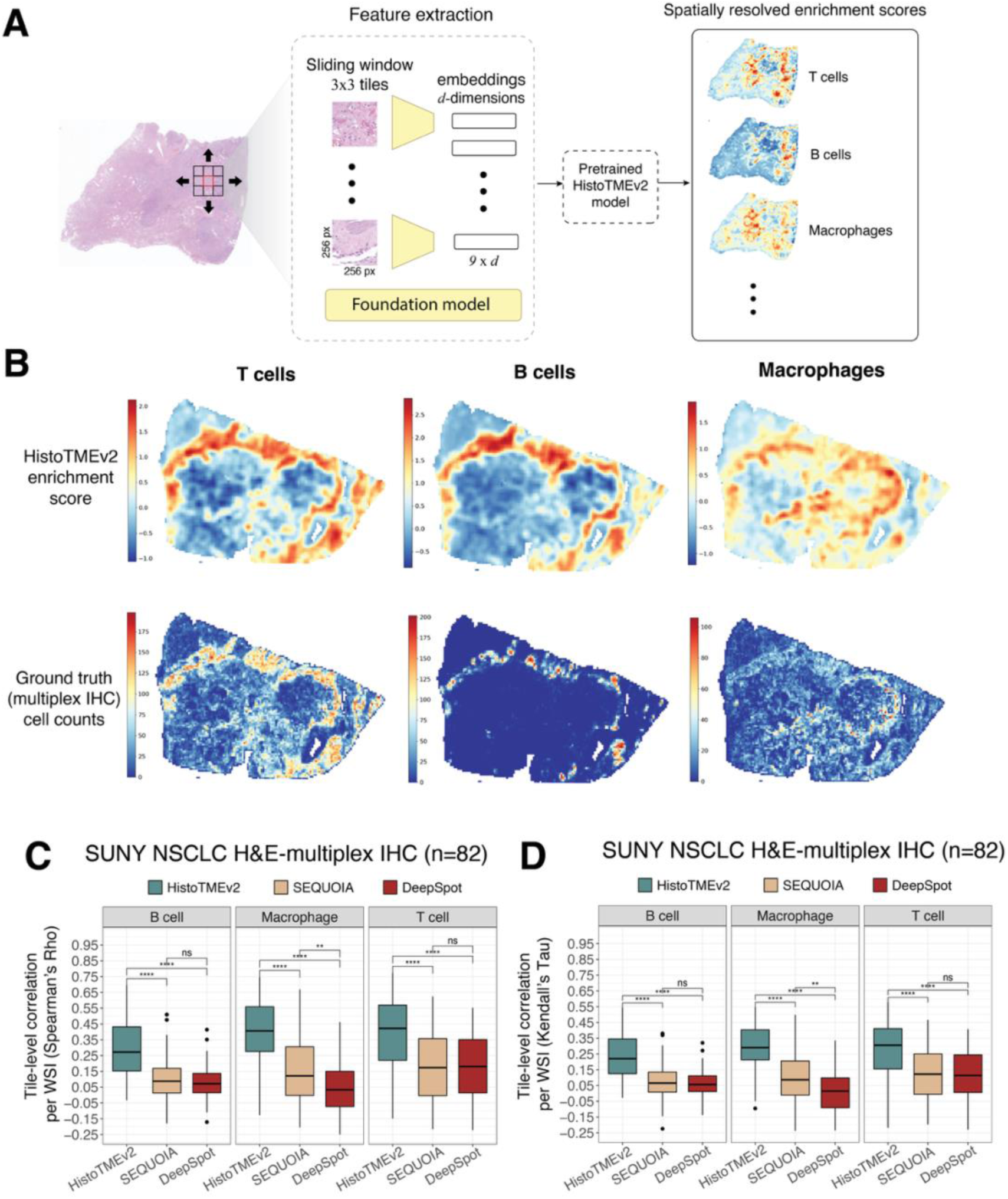
Spatial prediction performance of HistoTME-v2 on SUNY NSCLC cohort with spatially registered multiplex IHC. **(A)** Schematic overview of spatial inference using HistoTME-v2, utilizing a sliding window context of 3x3 tiles **(B)** Representative non-small cell lung cancer (NSCLC) sample from the SUNY Upstate cohort, showing spatial enrichment scores predicted by HistoTME-v2 for T cells, B cells, and macrophages, alongside spatially matched multiplex IHC-based measurements for the same cell types. **(C)** Tile-level (∼112×112 µm) Spearman correlations between predicted enrichment scores for T cells, B cells, and macrophages and observed single-cell counts from spatially registered multiplex IHC sections in the SUNY Upstate NSCLC cohort. **(D)** Tile-level (∼112×112 µm) Kendall’s Tau correlations between predicted enrichment scores for T cells, B cells, and macrophages and observed single-cell counts from spatially registered multiplex IHC sections in the SUNY Upstate NSCLC cohort.

Interestingly, although HistoTME-v2 was never trained or finetuned on spatial transcriptomics datasets, it significantly outperformed both the baseline models, SEQUOIA and DeepSpot in capturing spatial distribution of T cells, B cells and Macrophages in the lung TME (See **Table 1** and **Figure 3C**).

**Table 1.** Spearman correlation between predicted cell type-specific enrichment scores and observed single-cell counts per tile across models. HistoTME-v2 was compared against SEQUOIA and DeepSpot using paired H&E and multiplex IHC data from the SUNY Upstate NSCLC cohort.

| Model | T Cells (Median [Min, Max]) | Macrophages (Median [Min, Max]) | B Cells (Median [Min, Max]) |
| --- | --- | --- | --- |
| **HistoTME-v2** | 0.45 [-0.28, 0.82] | 0.41 [-0.12, 0.75] | 0.29 [-0.03, 0.81] |
| **SEQUOIA** | 0.17 [-0.30, 0.71] | 0.12 [-0.32, 0.67] | 0.08 [-0.29, 0.57] |
| **DeepSpot** | 0.17 [-0.40, 0.61] | 0.014 [-0.55, 0.46] | 0.075 [-0.31, 0.72] |

Given that certain immune cell types—such as B cells—may be sparsely distributed within the TME^47^, we also calculated Kendall’s Tau correlations as a robustness check. These results were consistent with the Spearman-based trends (**Table 2**, **Figure 3D**). A notable trend across models was that samples with higher abundances of a given cell type exhibited greater spatial prediction accuracy for that cell type, while samples with lower abundances showed reduced performance (**Supplementary** Figure 1)—indicating that cell type sparsity imposes a fundamental limitation on accurate spatial modeling of the TME from H&E alone.

**Table 2.** Kendall’s Tau correlation between predicted cell type-specific enrichment scores and observed single-cell counts per tile across models. HistoTME-v2 was compared against SEQUOIA and DeepSpot using paired H&E and multiplex IHC data from the SUNY Upstate NSCLC cohort.

| Model | T Cells (Median<br>[Min, Max]) | Macrophages (Median<br>[Min, Max]) | B Cells (Median<br>[Min, Max]) |
| --- | --- | --- | --- |
| **HistoTME-v2** | 0.32 [-0.22, 0.62] | 0.29 [-0.1, 0.55] | 0.23 [-0.02, 0.62] |
| **SEQUOIA** | 0.12 [-0.2, 0.51] | 0.09 [-0.24, 0.5] | 0.07 [-0.22, 0.41] |
| **DeepSpot** | 0.12 [-0.29, 0.41] | 0.01 [-0.34, 0.34] | 0.06 [-0.25, 0.55] |

**Table 3:** Summary of Cohorts Used for Training and External Validation of HistoTME-v2.

| Cohort Name | Validation Level | # Patients / Samples | Tumor Types | Data Modalities |
| --- | --- | --- | --- | --- |
| **TCGA**<sup>33</sup> | Training | 6,901 patients / 7,586 WSIs | 24 cancer types | H&E-stained whole slide images (WSIs), bulk RNA sequencing |
| **CPTAC**<sup>34</sup> | External – Patient/WSI-Level | 1,693 patients / 5,433 WSIs | 7 cancer types | H&E-stained WSIs, bulk RNA sequencing |
| **NCI Prostate cancer**<sup>36</sup> | External – Patient/WSI-Level | 36 patients / 486 treatment naïve biopsy images | Advanced prostate cancer | H&E-stained biopsies, bulk RNA sequencing |
| **Fred Hutch HGSOC**<sup>35</sup> | External – Patient/WSI-Level | 46 patients / 92 WSIs | High-grade serous ovarian cancer | H&E-stained WSIs, bulk RNA sequencing |
| **SUNY Upstate NSCLC**<sup>32</sup> | External – WSI, Tile, IHC-Level | 82 patients / 254 WSIs + 82 IHC | Non-small cell lung cancer (NSCLC) | H&E-stained WSIs, spatially registered multiplex IHC (CD3, CD4, CD8, CD20, CD163) |
| **CRC-FFPE-CODEX**<sup>45</sup> | External – Region-Level | 35 patients / 140 TMA cores | Advanced colorectal cancer | H&E-stained TMA slides, CODEX multiplex imaging (56 protein markers) |
| **HEST1K FFPE-Visium**<sup>48</sup> | External – Spot-Level | 37 WSIs (selected from a total of 354 tumor WSI) | 7 cancer types | H&E-stained WSIs, Visium spatial transcriptomics (UMI > 10,000 per spot) |

**Table 4:** Summary of Evaluation Metrics Used for External Validation of HistoTME-v2 Across different Ground Truth Data Modalities and Spatial Scales. This table outlines the performance metrics employed to validate model predictions against diverse ground truth data types. For datasets with matched bulk transcriptomics, Pearson correlation was used to quantify agreement between predicted and observed signature scores. For single-cell resolved cell type proportions derived from multiplex IHC or CODEX, Spearman rank correlation was applied to accommodate potential non-linearities and scale differences. For spatially resolved evaluations, Kendall’s Tau was additionally included—alongside Pearson and Spearman—to address the impact of sparsity in spatial distributions of certain immune cell types such as B cells.

| Data Type | Ground Truth Modality | Metric(s) Used | Rationale |
| --- | --- | --- | --- |
| **WSI with bulk transcriptomics** | Bulk RNA Seq | Pearson correlation | Assesses linear correlation between predicted and bulk-derived signature scores |
| **WSI with single-cell cell type fractions** | Multiplex IHC or CODEX | Spearman rank correlation | Captures rank-based relationships and accommodates scale differences and nonlinearity |
| **WSI with spatially resolved cell type/signature distributions** | Multiplex IHC/CODEX/Visium (spatial transcriptomics) | Pearson, Spearman, Kendall's Tau | Besides Pearson and Spearman, Kendall's Tau is included to better handle sparse and noisy spatial distributions (e.g., B cells) |

Second, we benchmarked the spatial prediction performance of HistoTME-v2 on the HEST- 1K database^48^, a curated collection of 1000 whole slide H&E images and spatial transcriptomic datasets designed to support benchmarking of computational methods for spatial gene expression prediction from whole slide histopathology images. From this database, we specifically focused on validation on FFPE-Visium samples due to their broad coverage of the transcriptome, allowing us to systematically evaluate all predicted transcriptomic signatures. However, given known technical limitations of Visium in archival FFPE tissue—such as mRNA degradation and frequent gene dropouts^13^—we restricted our benchmarking analysis to FFPE-Visium slides with a median of at least 10,000 UMIs captured per spot to ensure meaningful comparisons^49^. This initial filtering resulted in a reduced cohort of 37 high-quality FFPE-Visium samples from HEST-1K spanning 7 different cancer types (See Methods). In this benchmarking experiment, HistoTME-v2 was primarily compared with SEQUOIA. DeepSpot was excluded as it was previously trained on HEST-1K data^46^.

Across all samples evaluated, HistoTME-v2 most accurately predicted the spatial distribution of extracellular matrix and cancer-associated fibroblast (CAF) signatures, achieving, high spot-level correlation coefficients per WSI (median 0.68 and 0.69, respectively). These significantly outperformed the baseline model, SEQUOIA’s predictions (median 0.48 and 0.53; p < 0.001; see **Figure 4A**, **Figure 4B**). The higher accuracies achieved for stromal signatures compared to immune signatures by both models reflects the higher spatial variability and detection rates of stromal genes compared to immune-related genes in Visium samples (**See Supplementary** Figure 2). Despite these challenges, HistoTME-v2 consistently outperformed the baseline across almost all evaluated transcriptomic signatures.

**Figure 4:**
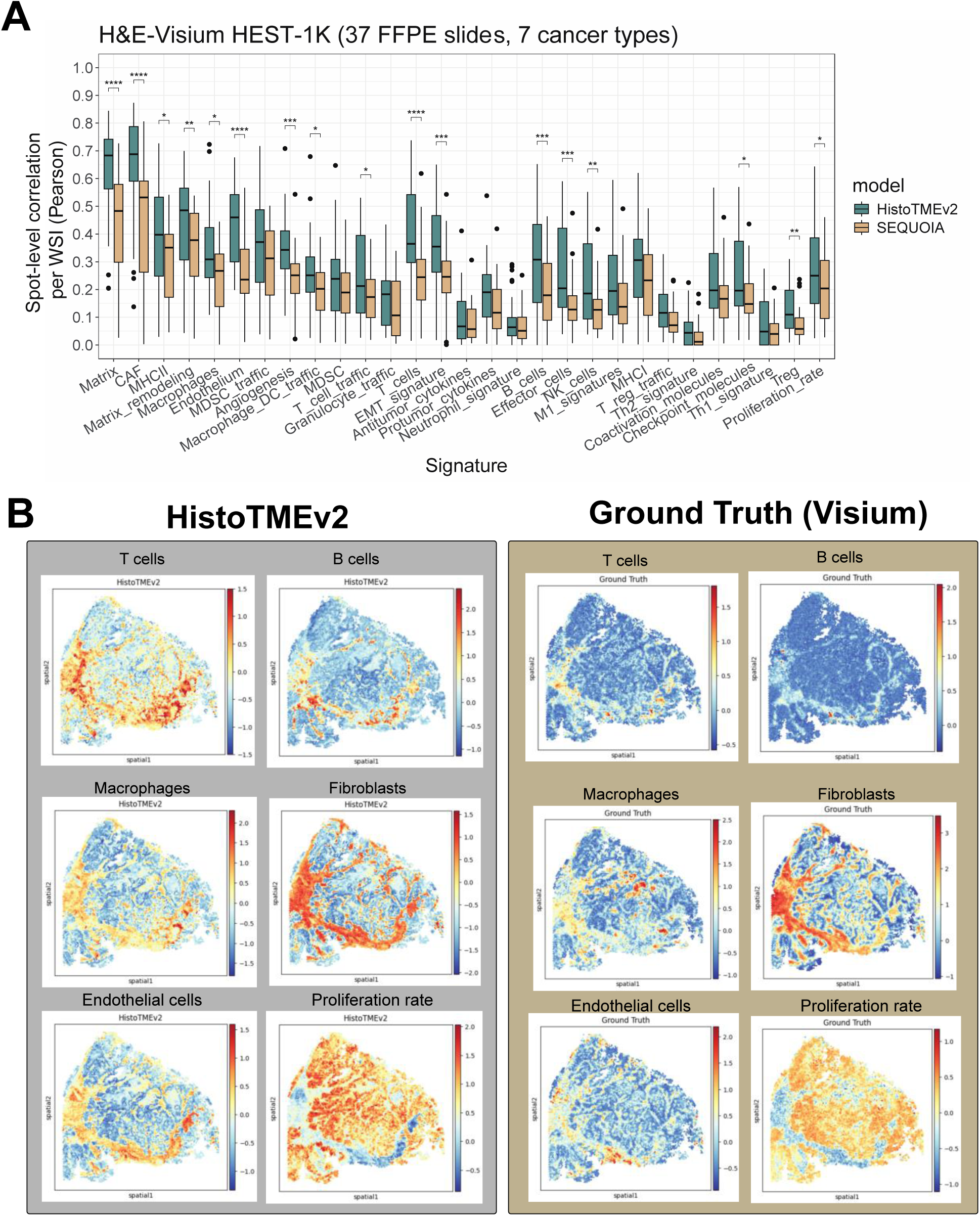
Spatial prediction performance of HistoTME-v2 on HEST1K FFPE-Visium data. **(A)** Spot-level (∼55 µm) correlation of predicted enrichment scores with Visium-derived enrichment scores across selected high-quality Visium samples (median >10,000 UMIs/spot) from the HEST1K database. Statistical comparisons were performed using the paired Wilcoxon test. CRC: Colorectal cancer; NSCLC: Non-small cell lung cancer. **(B)** Representative example of Spot-level (∼55 µm) enrichment scores predicted by HistoTME-v2 for T cells, B cells, macrophages, fibroblasts, endothelial and malignant cell components across a high-quality Visium sample (median 49885 UMIs/spot) from the HEST1K database, compared against ground truth Visium-derived gene signature scores.

Taken together, these results demonstrate the enhanced robustness of HistoTME-v2 for characterizing the spatial organization of diverse TME components across multiple tissue types.

## Discussion

In summary, HistoTME-v2 enables in-depth molecular and spatial characterization of the TME directly from standard H&E-images—without requiring nuclear segmentation, cell classification, or generation of virtual stains. To our knowledge, this is the first study to rigorously validate such a framework across 25 distinct cancer types and different platforms. In contrast to several recent methods aiming to reconstruct whole bulk or spatial transcriptomes from H&E images^31,38,47,50^, HistoTME-v2 focuses on modeling expression patterns of curated biologically interpretable gene sets. Our extensive benchmarking experiments across multiple cancer types demonstrate that this approach often yields more robust and accurate estimates of the TME composition compared to those derived from imputed gene expression profiles. This aligns well with findings from several single-cell studies emphasizing the stability and functional relevance of transcriptional signatures— which often offer more meaningful biological insights compared to noisy single-gene measurements^51,52^. Moreover, our spatial benchmarking results indicate that HistoTME-v2 outperforms models explicitly trained on spatially matched H&E and spatial transcriptomic data, generating more reliable spatial predictions. This further highlights the generalizability and effectiveness of leveraging weak supervision and foundation model-based representations for histology-driven spatial profiling.

While HistoTME-v2 focuses on 29 curated TME-related transcriptomic signatures, the current framework can be further extended to characterize additional transcriptional signatures directly from H&E slides. For example, HistoTME-v2 could be further expanded to precisely characterize cellular plasticity^53^ and hypoxia-associated signatures^54,55^. Another promising application is to characterize signatures associated with tertiary lymphoid structure formation and function^56,57^, an important factor influencing immunotherapy outcomes in multiple cancer types^58^.

With its significantly expanded scope, HistoTME-v2 can now be leveraged to investigate the TME in a broad range of cancer types and clinical contexts using only H&E-stained pathology slides. We previously demonstrated the clinical utility of such an approach in advanced non- small cell lung cancer, where we retrospectively identified additional immunotherapy responders missed by conventional PD-L1 IHC scoring^32^. One promising new direction for HistoTME-v2 is its application to high risk (Gleason >= 4+4) or metastatic hormone-sensitive prostate cancer, particularly in the analyzing tumor-adjacent stroma—an area with a critical unmet need for biomarkers to guide early intervention and delay castration resistance^59,60^. Recent single-cell and spatial transcriptomic studies have revealed substantial heterogeneity in the TME composition of prostate cancer across disease stages, offering insights into novel mechanisms driving disease progression^36,60–63^. Hence, leveraging HistoTME-v2 to profile the TME of advanced prostate cancer directly from diagnostic H&E slides could offer a scalable and cost-effective avenue for discovering additional biomarkers, ultimately advancing personalized treatment strategies. Another promising direction is the study of spatio-temporal TME dynamics using H&E image data sampled from tumor organoids^64^.

Despite its advantages, HistoTME-v2 has certain limitations that can be improved upon in future work. Notably, it does not produce predictions at single-cell resolution. Instead, its spatial outputs are constrained by sliding window predictions over 3×3 tile patches, which capture a broader tissue context. This coarser resolution limits the ability of HistoTME-v2 to quantify local cell–cell interactions. However, this design trade-off allows for efficient, whole-tissue spatial analysis without the substantial computational overhead of segmenting and analyzing millions of cells. Future work should explore further refining HistoTME-v2’s spatial predictions by integrating single-cell segmentation tools—such as CellVIT^15^—thereby enabling single-cell spatial analysis of the tumor microenvironment from H&E-stained images.

Second, HistoTME-v2 was primarily trained on diagnostic FFPE specimens paired with bulk transcriptomics, owing to their widespread availability and higher image quality compared to fresh frozen (FF) specimens^65^. As such, its performance on FF tissues has not been evaluated. Future studies should explore incorporating FF data during training and validation to further improve model generalizability.

Third, while we externally validated HistoTME-v2 across diverse institutional datasets, we did not systematically assess robustness of predictions to variations in image scanning quality. Recent studies have shown that foundation model embeddings may capture institution-specific biases, potentially confounding downstream applications and generalizability^26,66^. To mitigate specific foundation model biases, we took an ensemble approach, which on average showed improved robustness and accuracy over any single foundation model. However, additional benchmarking and stress-testing under varying conditions may be needed to further strengthen robustness.

Lastly, although the predicted enrichment scores show strong correlation with single-cell abundances, they should be interpreted as relative, rather than absolute measures of cell type abundance. We also observe an inherent limitation in predicting the spatial distribution of cell types, which appears to be constrained by their overall abundance (**Supplementary** Figure 1)—suggesting that rare cell types or specialized subsets may be particularly challenging to infer from H&E images alone and may require more advanced imaging techniques. Future work should investigate cancer-type-specific calibration and fine-tuning of HistoTME-v2 predictions to more precisely map the relationship between predicted enrichment scores and actual single-cell densities across diverse tumor contexts.

Overall, by linking subtle morphological variations in tissue with biologically interpretable transcriptomic signatures, HistoTME-v2 presents a powerful new tool to study and model the TME from large-scale digital pathology image repositories, further advancing biomarker discovery and the mission of precision oncology.

## Code availability

https://github.com/spatkar94/HistoTME/tree/main

## Funding Statement & Acknowledgements

This project was supported by an award from Upstate Foundation’s Hendricks Endowment as well as intramural funding from the National Cancer Institute, NIH. Data (digital images and clinical meta-data) from the SUNY institutional cohort was generated at the Pathology Research Core Lab using institutional resources and support.

## Methods

### Datasets

The training and validation of HistoTME-v2 were performed using a diverse collection of datasets spanning multiple tumor types, imaging modalities, and transcriptomic platforms.

- **TCGA (Training and internal validation):** The primary training dataset consisted of 7,586 formalin-fixed paraffin-embedded (FFPE), H&E-stained whole slide images (WSIs) paired with processed bulk RNA sequencing data from 6,901 patients, representing 24 distinct tumor types from The Cancer Genome Atlas Network (TCGA^33^). All data are publicly available and obtained from the Genomic Data Commons (GDC) portal (https://portal.gdc.cancer.gov/).
- **CPTAC-3 (External Validation – Patient/Whole Slide-Level):** A total of 5,433 FFPE H&E-stained WSIs from 1,693 patients were obtained from the Clinical Proteomic Tumor Analysis Consortium (CPTAC-3^34^; https://proteomics.cancer.gov/programs/cptac), covering seven different cancer types. Each sample included matched bulk RNA sequencing data. To avoid data leakage, only patients with no overlap with TCGA were included. CPTAC WSI data was obtained from the Cancer Imaging Archive (TCIA; https://www.cancerimagingarchive.net/) and the corresponding transcriptomics data was obtained from the genomic data commons
- **NCI Prostate Cancer (External Validation – Patient/Whole Slide-Level):** 486 H&E-stained treatment-naïve tumor biopsy images from 36 patients with advanced prostate cancer enrolled in a neoadjuvant ADT + Enzalutamide trial (NCT02430480) were utilized for external validation. Each biopsy was matched with bulk RNA sequencing derived from the same tumor block (GEO accession: GSE183100^36^). The imaging data are available from TCIA (https://doi.org/10.7937/TCIA.JHQD-FR46)
- **Fred Hutchinson HGSOC Cohort (External Validation – Patient/Whole Slide- Level):** 92 FFPE H&E-stained WSIs from 46 patients with high-grade serous ovarian cancer (HGSOC) were downloaded from a recent study conducted at the Fred Hutchinson Cancer Center^35^. Each slide was accompanied by matched processed bulk transcriptomic data made available by the original study authors. The WSI data are available for download from TCIA (https://doi.org/10.7937/6RDA-P940)
- **SUNY Upstate NSCLC Cohort (External Validation – Patient/Whole slide level, Tile-Level):** This cohort included 254 FFPE H&E-stained surgical tissue slides from 82 patients with non-small cell lung cancer (NSCLC), along with adjacent multiplex IHC slides stained for T cells (CD3+, CD4+, CD8+), B cells (CD20+), and macrophages (CD163+). To estimate single cell fractions (% marker positive cells) from IHC, tumor regions were manually segmented, and immune cells were quantified using in-built stain deconvolution and positive cell detection algorithm implemented in QuPath v0.5.0(https://qupath.readthedocs.io/en/stable/docs/tutorials/cell_detection.html). To facilitate spatial validations, each IHC image was spatially registered to its corresponding source H&E image using the Vallis 1.1.0 spatial registration package^67^. This institutional dataset was approved for research use by the SUNY Upstate Institutional Review Board (IRB #1857564, #2153970) and considered exempt from further IRB oversight. All data were collected retrospectively and anonymized. Additional details on slide preparation and scanner are available in our prior publication^32^.
- **CRC-FFPE-CODEX Cohort (External Validation – Region-Level):** A total of 140 tissue microarray (TMA) cores were collected from the invasive tumor fronts of 35 patients with advanced colorectal cancer45. Each TMA core was matched with FFPE H&E-stained slides and corresponding CODEX multiplex imaging data quantifying 56 protein markers. Single-cell fractions (%marker positive cells) of various immune, stromal, endothelial, and tumor cell populations within each core were previously quantified by the original study authors from raw CODEX images and made publicly available (https://data.mendeley.com/datasets/mpjzbtfgfr/1). For benchmarking purposes, six major cell types—T cells, B cells, macrophages, fibroblasts, endothelial cells, and Ki67⁺ (proliferating) cells—were selected based on their ability to be reliably matched with corresponding HistoTME-v2-derived cell type–specific signature scores. In cases where multiple immune subsets (e.g., CD4⁺, CD8⁺, and CD3⁺ T cells) were reported, their counts were aggregated to enable one-to-one comparison with broader cell-type–level predictions from HistoTME-v2. All raw H&E and CODEX imaging data used in this study are available for download from The Cancer Imaging Archive (TCIA) (https://doi.org/10.7937/TCIA.2020.FQN0-0326)
- **HEST-1K FFPE-Visium (External Validation – Spot-Level):** The HEST1K collection^48^ consists of 1000 matched H&E WSIs and spatial transcriptomics profiles, spanning both Visium and Xenium platforms. Given the variability in image and spatial transcriptomics data quality within HEST1K, a three-step filtering procedure was applied prior to downstream benchmarking. First, only FFPE-Visium tumor samples with a median UMI count per spot >10,000 were retained to ensure high signal-to-noise ratios and comprehensive spatial benchmarking of all predicted signatures. This resulted in a high-quality cohort of 37 FFPE-Visium tumor samples spanning 7 cancer types: Colorectal carcinoma (n=21), renal cell carcinoma (n=8), Prostate Adenocarcinoma (n=2), Serous Ovarian Carcinoma (n=2), Lung-Squamous Cell Carcinoma (n=1), Lung-Neuroendocrine Carcinoma (n=1), Invasive Ductal Carcinoma (n=1), Cervical Squamous Cell Carcinoma (n=1). Second, spots with <1,000 UMIs were filtered out and third genes with <100 counts per spot were filtered out to remove low-abundance and potentially uninformative transcripts. The remaining UMI count data were normalized by dividing by the total counts per spot and used in downstream benchmarking analyses.

### WSI Preprocessing

Each WSI underwent tissue segmentation and was tessellated into 256×256 pixel tiles at 20x magnification (∼112×112 μm), using the Trident whole slide imaging preprocessing package^68^. This resolution was selected to capture both local nuclear features and the surrounding tissue context, while also facilitating systematic benchmarking against other published models that operate at comparable scales. Tiles were encoded into feature embeddings using six foundation models: UNI_v1^39^, UNI_v2^68^, Virchow^40^, Virchow2^41^, Hoptimus0^43^, and Gigapath^42^. These generated a bag of embeddings of size *n x d*, where n is the number of tiles and d is the model-specific embedding dimension. For TCGA slides, tiles were additionally stain-normalized using the Macenko method^69^ to avoid overfitting to staining variations across different cancer types and centers during training.

### Baseline Models

The primary baseline model used for pan-cancer benchmarking of HistoTME-v2 is SEQUOIA^38^, a linearized transformer architecture developed to predict bulk transcriptomic profiles directly from H&E-stained whole slide images (WSIs). SEQUOIA was trained on 7,584 tumor samples spanning 16 cancer types from The Cancer Genome Atlas (TCGA) and demonstrated strong generalization performance when validated on two independent external cohorts comprising 1,368 tumors. SEQUOIA is trained separately for each cancer type resulting in 16 cancer type-specific pretrained model weights. The full implementation of SEQUOIA is publicly available at https://github.com/gevaertlab/sequoia-pub, with pretrained model weights for each of the 16 cancer types accessible via https://huggingface.co/gevaertlab. Beyond bulk transcriptomic inference, SEQUOIA also supports spatially resolved gene expression prediction at loco-regional levels within tumor slides utilizing a sliding window approach with a user-specified window size, enabling localized analysis of tumor microenvironment heterogeneity. Given publicly accessible model weights, ease of implementation, and previously reported state of the art performance in pan cancer gene expression prediction from WSIs, SEQUOIA was selected as a strong baseline comparator for benchmarking HistoTME-v2.

Besides SEQUOIA, we also evaluated DeepSpot^46^, a recent state-of-the art deep learning model harnessing digital pathology foundation models and multiple-instance learning architectures to predict spatial gene expression from individual H&E image patches. Specifically, DeepSpot leverages foundation model embeddings extracted from a H&E image patch spatially aligned with a Visium spot, its neighboring patches as well as sub-patches and aggregates them using a set-based multiple instance learning architecture to predict the expression of all genes in that spot. DeepSpot has been trained on paired H&E and Visium spatial transcriptomics datasets from the publicly available HEST-1K database and private institutional data from the University Hospital of Zurich spanning lung, kidney and colon cancer types. Publicly available pretrained model checkpoints for DeepSpot are available at (https://zenodo.org/records/15322099).

### Computing Signature Enrichment Scores from Bulk and Spatial Transcriptomics

Normalized bulk RNA-seq data (gene-level TPM counts) from TCGA, CPTAC, NCI, and Fred Hutch cohorts were downloaded from public sources (see Table 1) and log₂-transformed. Following this, sample specific gene set enrichment scores were calculated based on ranked expression of 29 functional gene sets previously defined by Bagaev et al.^37^, thereby comprehensively capturing the TME cellular and molecular composition. The same pipeline was applied for computing enrichment scores from H&E-imputed transcriptomic data. To compute gene set enrichment scores from spatial transcriptomics data, the *sc.tl.score_genes* function in Scanpy^70^ was used, which follows a similar signature scoring algorithm as the one implemented in Bagaev et al^71^.

### Pan-Cancer Model Training and Evaluation

HistoTME-v2 uses the same architecture as the original HistoTME model^32^, comprising a foundation model backbone and a shared task-specific ABMIL module for weakly supervised regression. The output is a single vector of gene set enrichment scores,one per transcriptomic signature. Unlike HistoTME, which was trained solely on NSCLC, HistoTME- v2 was trained on 7,586 WSIs with matched transcriptomic data across multiple tumor types from TCGA.

Model training followed a 5-fold cross-validation protocol using stratified K-fold splits (80% training, 20% validation) using the scikit-learn package. Optimization was performed using the AdamW optimizer and the Huber loss function (δ = 1):

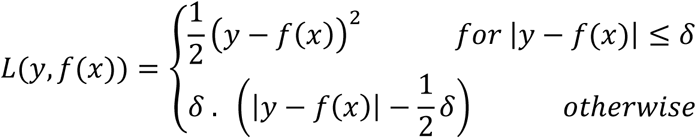

The learning rate and weight decay were both set to 1x 10-4. Due to the variable size of input bags, a batch size of 1 was used with gradient accumulation over 8 steps. Each foundation model variant was trained for up to 40 epochs, with early stopping applied if validation performance did not improve for 3 consecutive epochs. Model performance was evaluated using Pearson correlation on a held-out validation set. An ensemble model, created by averaging predictions from all foundation models, achieved the highest mean correlation across cross-validation folds and was therefore selected for external validation. All models were implemented in Python using PyTorch v2.7.0 and trained on a server equipped with four NVIDIA A100 GPUs.

### Performance Evaluation Metrics

Several performance evaluation metrics were considered in this study, tailored to the varying spatial resolutions and ground truth data modalities (bulk/spatial transcriptomics, multiplex IHC, CODEX) available for external validations. For WSI datasets with matched bulk transcriptomics data, we utilized the Pearson correlation metric, measuring the correlation between AI-predicted signature scores and bulk transcriptomics–derived signature scores. For WSI datasets with ground truth single-cell fractions of specific cell types, measured from either multiplex IHC or CODEX, we utilized the Spearman rank correlation metric to evaluate accuracy, considering the different scales of measurement and nonlinear relationships between predicted scores and actual cell counts. At the spatial level, the Kendall’s Tau metric was calculated in addition to Pearson and Spearman Correlation Metric to account for high sparsity in spatial distribution measurements of certain immune cell types (e.g., B cells).

## Supplementary Figures

**Supplementary Figure 1:**
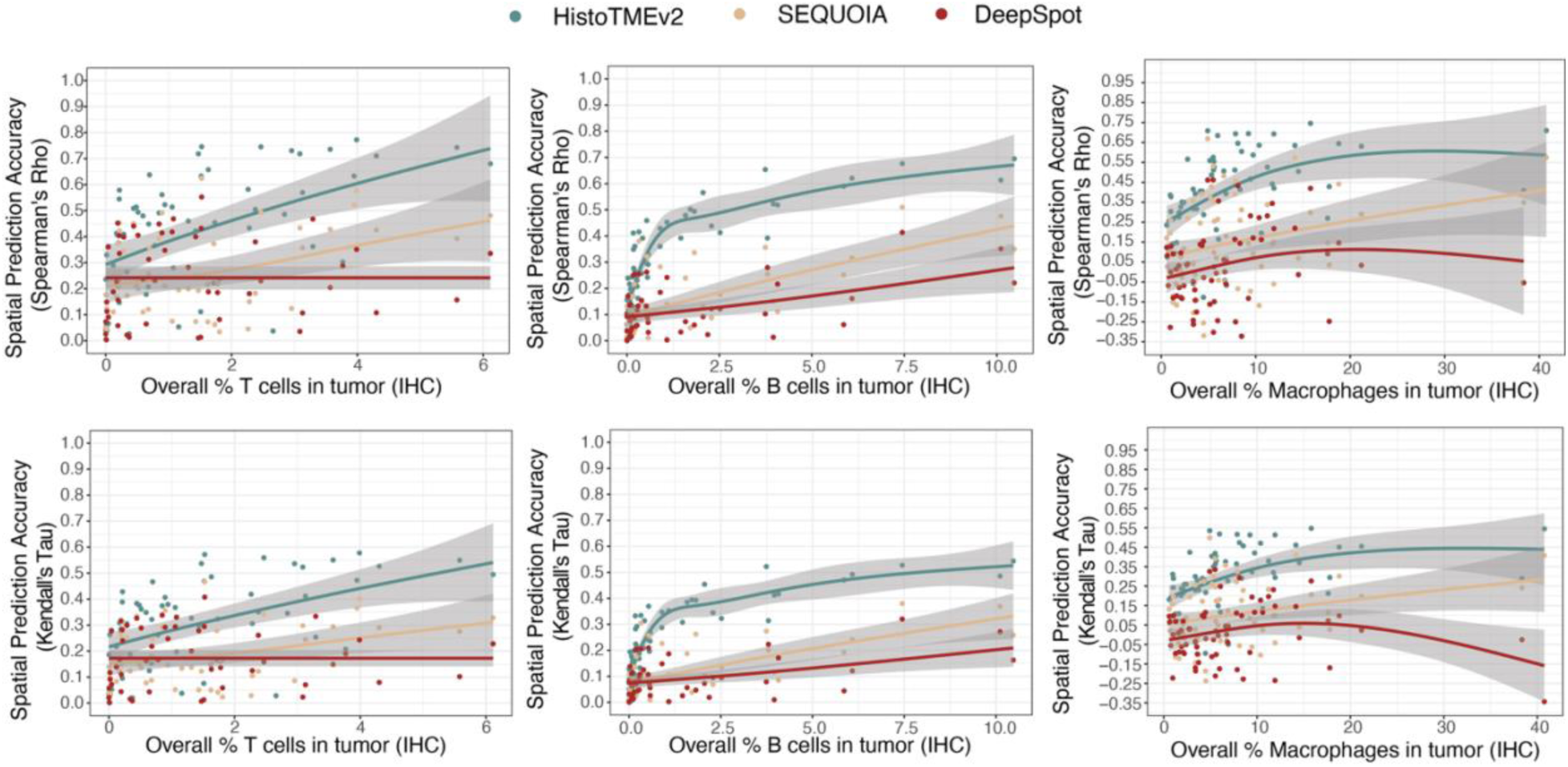
Relationship between spatial prediction performance and cell type abundance. Top three panels show scatterplots illustrating the association between spatial prediction performance (Spearman correlation) and the abundance of each immune cell type (measured as % marker-positive cells per sample). A LOESS regression line was fitted to highlight trends. Bottom three panels present the same analysis using the Kendall’s Tau correlation metric. Data shown is from the SUNY NSCLC cohort.

**Supplementary Figure 2:**
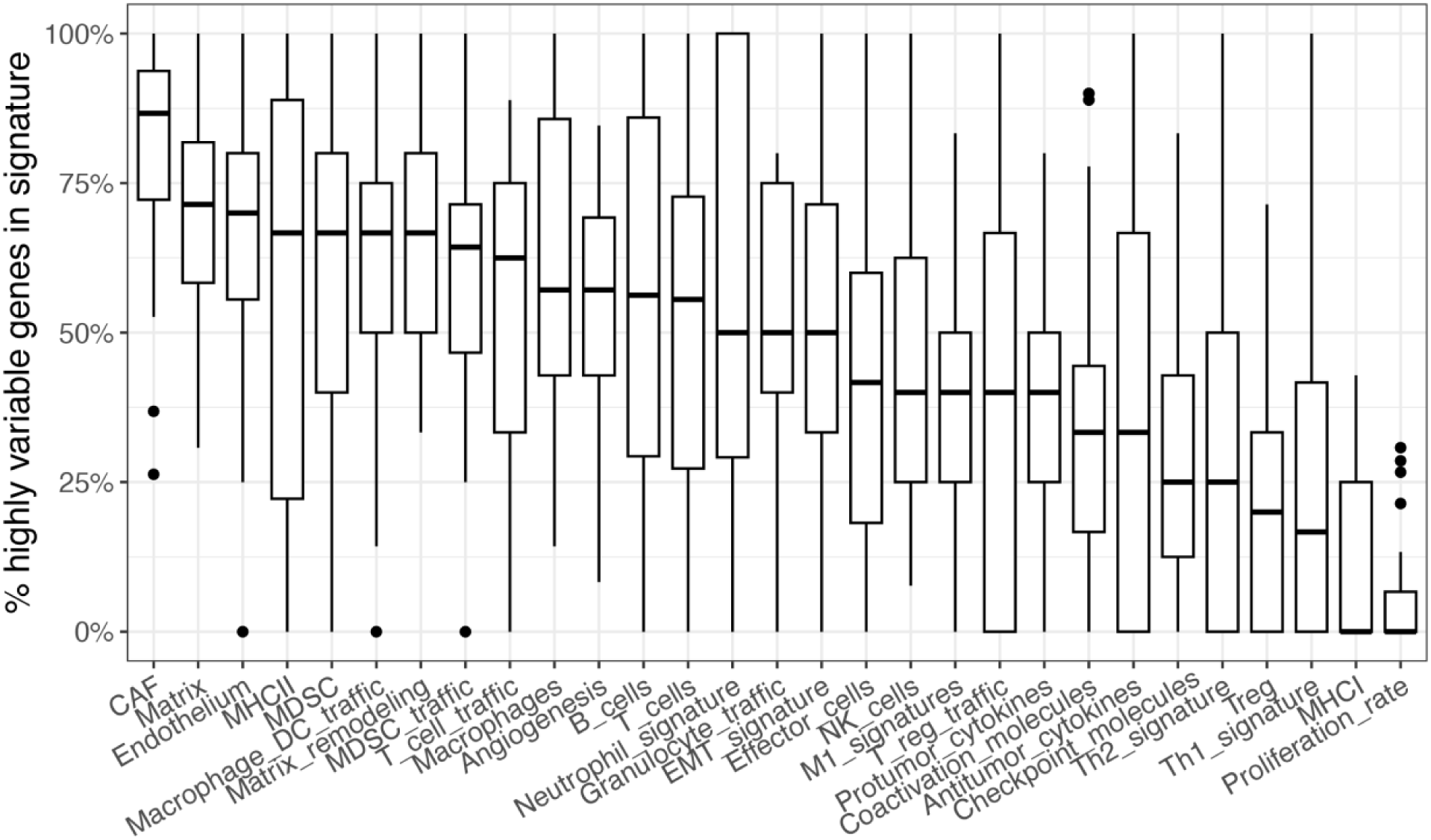
Proportion of highly spatially variable genes within each transcriptomic signature in the HEST1K FFPE-Visium cohort. Box plots show the distribution of the percentage of genes classified as highly spatially variable within each signature across whole-slide images (WSIs). Spatial variability was determined per WSI using spatial transcriptomics data and the `scanpy.pp.highly_variable_genes` function with the Seurat v3 flavor72.

