## Supplementary figures and images for "Towards interpretable molecular and spatial analysis of the tumor microenvironment from digital histopathology images with HistoTME-v2"

### Supplementary Figure 1

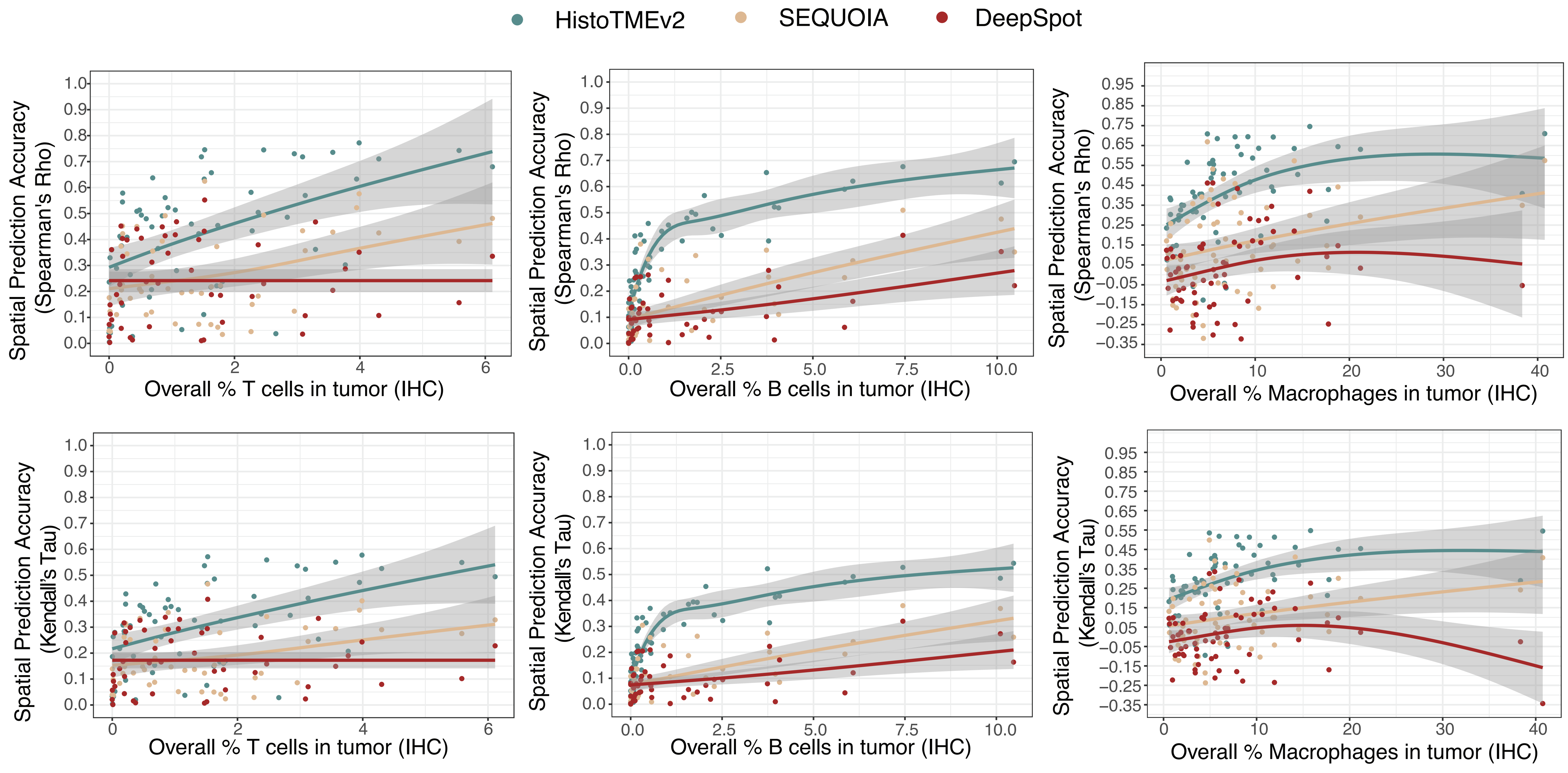

### Supplementary Figure 2

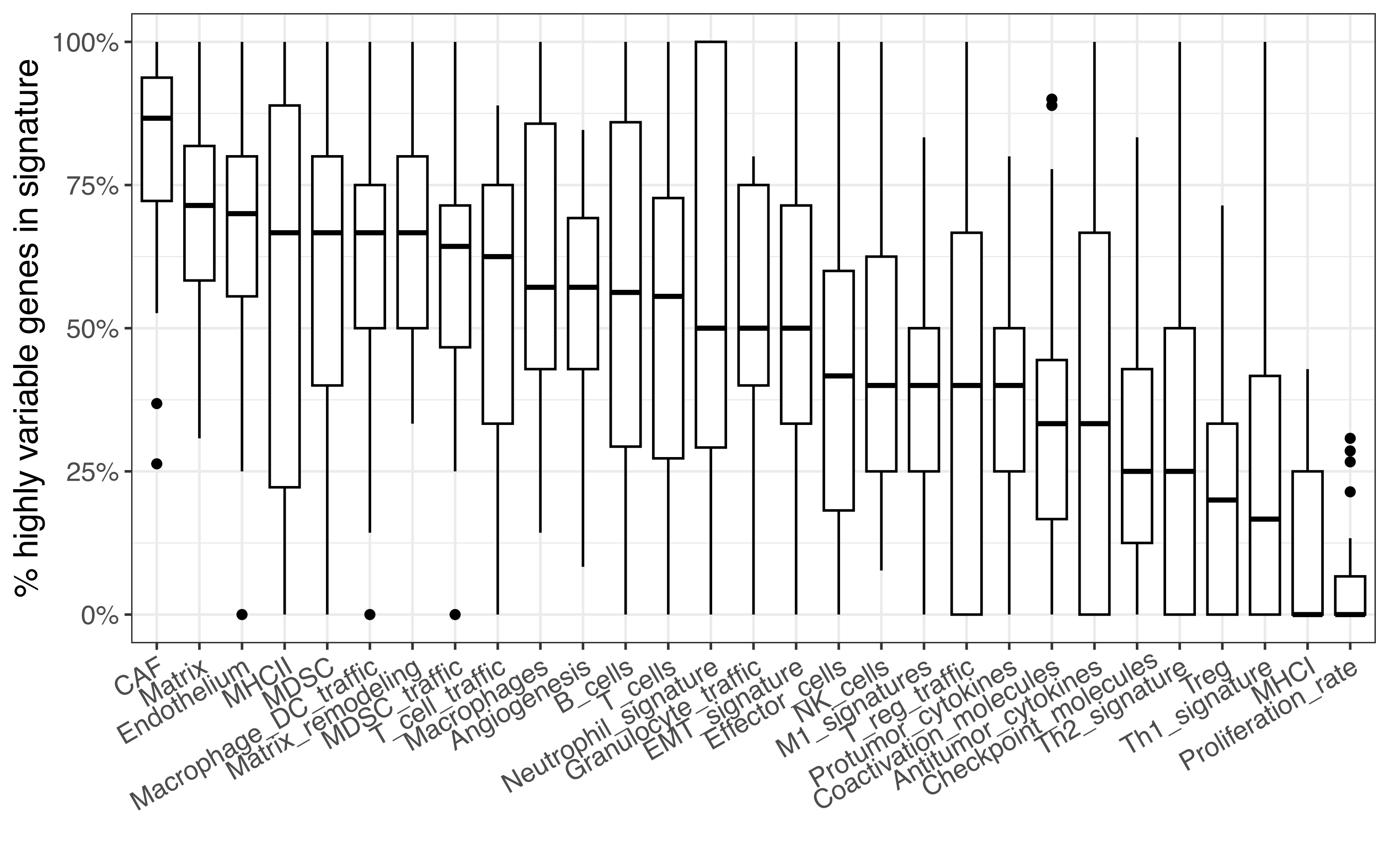
